# Short homology-directed repair using optimized Cas9 in the pathogen *Cryptococcus neoformans* enables rapid gene deletion and tagging

**DOI:** 10.1101/2021.06.30.450566

**Authors:** Manning Y. Huang, Meenakshi B. Joshi, Michael J. Boucher, Sujin Lee, Liza C. Loza, Elizabeth A. Gaylord, Tamara L. Doering, Hiten D. Madhani

**Author notes:** co-first authors (Manning Y. Huang, Meenakshi B. Joshi).

## Abstract

*Cryptococcus neoformans*, the most common cause of fungal meningitis, is a basidiomycete haploid budding yeast with a complete sexual cycle. Genome modification by homologous recombination is feasible using biolistic transformation and long homology arms, but the method is arduous and unreliable. Recently, multiple groups have reported the use of CRISPR-Cas9 as an alternative to biolistics, but long homology arms are still necessary, limiting the utility of this method. Since the *S. pyogenes* Cas9 derivatives used in prior studies were not optimized for expression in *C. neoformans*, we designed, synthesized, and tested a fully *C. neoformans*-optimized Cas9. We found that a Cas9 harboring only common *C. neoformans* codons and a consensus *C. neoformans* intron together with a *TEF1* promoter and terminator and a nuclear localization signal (*C. neoformans*-optimized *CAS9* or “*CnoCAS9*”) reliably enabled genome editing in the widely-used KN99α *C. neoformans* strain. Furthermore, editing was accomplished using donors harboring short (50 bp) homology arms attached to marker DNAs produced with synthetic oligonucleotides and PCR amplification. We also demonstrated that prior stable integration of *CnoCAS9* further enhances both transformation and homologous recombination efficiency; importantly, this manipulation does not impact virulence in animals. We also implemented a universal tagging module harboring a codon-optimized fluorescent protein (mNeonGreen) and a tandem Calmodulin Binding Peptide-2X FLAG Tag that allows for both localization and purification studies of proteins for which the corresponding genes are modified by short homology-directed recombination. These tools enable short-homology genome engineering in *C. neoformans*.

## INTRODUCTION

The opportunistic pathogenic yeast *Cryptococcus neoformans* is the most common cause of fungal meningitis and is responsible for ~15% of deaths in AIDS patients, causing ~200,000 deaths annually (Rajasingham *et al.* 2017). *C. neoformans* is a basidiomycete yeast and thus highly diverged from the ascomycete model yeasts *S. cerevisiae* and *S. pombe* (Janbon *et al.* 2014). Originally classified as a single species with four serotypes (A-D), these serotype designations continue to evolve (Hagen *et al.* 2017), with the most common serotype studied (serotype A) now called *C. neoformans* with seven pathogenic species recognized. As a model organism, it has many advantages including ease of cultivation, stable haploid cells, and a complete sexual cycle (Chun and Madhani 2010). Excellent animal models of infection have been developed, enabling studies of host-pathogen interactions (Cox *et al.* 2000). A congenic strain pair, KN99α and KN99**a**, has been developed and is widely used in the field (Nielsen *et al.* 2003).

Facile genetic manipulation is a cornerstone of the study of microbial pathogens, allowing the use of reverse genetics to dissect gene function, discover novel virulence determinants, and identify novel drug targets. Developed in the 1990s, biolistic transformation can achieve efficient editing in *C. neoformans* (Davidson et al. 2002). Indeed, our laboratory has used this method to construct a genome-scale gene knockout collection which we have made available through the Fungal Genetic Stock Center (www.fgsc.net). However, drawbacks of the biolistic method include a requirement for long (1kb) homology arms, an expensive biolistic instrument, costly gold nanoparticles, and other disposables. Moreover, biolistic transformation is finicky and not always reliable. Thus, there is an unmet need for a more facile method.

Several CRISPR-Cas9 systems have been developed for use in *C. neoformans*. In one system, biolistic delivery was used to introduce a self-processing sgRNA-encoding construct and a targeting construct harboring 1kb homology arms into a strain already carrying *CAS9* at the *SH1* locus (Arras *et al.* 2016). Another system used electroporation of a “suicide cassette” in which recombinational events excise CRISPR-Cas9 components from a cassette containing both a Cas9/sgRNA segment and a homology flanked-selectable marker (Wang *et al.* 2016). Both methods achieved high (70-90%) efficiencies of editing when targeting the *ADE2* locus. In the latter method, *CAS9* and sgRNA cassettes were lost in approximately half the resulting transformants, permitting later reuse of their method (Wang *et al.* 2016). A more recent method dubbed TRACE (<u>Tra</u>nsient <u>C</u>RISPR-Cas9 Coupled with <u>E</u>lecroporation) relies on transient expression of *CAS9* and sgRNAs from separate linear DNA molecules introduced by electroporation together with a homology donor DNA (Fan and Lin 2018). TRACE achieved *CAS9* and sgRNA cassette dose-dependent editing efficiencies between 40 to 90% when targeting the *ADE2* locus. Most recently, two additional transient methods using electroporation were described, one using a multi-guide approach, and a second using CRISPR-Cas9 RNPs to achieve high editing efficiency (Wang 2018).

Despite their utility, prior implementations of CRISPR-Cas9 in *C. neoformans* share several limitations. First, they require at least 500 bp arms for homology-directed recombination, which require potentially difficult construction by cloning or fusion PCR. Second, they were not optimized for Cas9 expression in *C. neoformans*. For example, the TRACE system uses a Cas9 derivative that was codon optimized for *C. elegans* and contains two *C. elegans* introns. Such introns would not be optimal for splicing in *C. neoformans* (Burke *et al.* 2018), although introns are necessary for gene expression in *C. neoformans* (Goebels *et al.* 2013). We reasoned that additional efficiency gains from codon optimization and use of an optimal intron might allow for shorter homology or improve CRISPR-Cas9 editing efficiency in this pathogen. We report here that a *Cryptococcus neoformans*-optimized Cas9 enables short homology-directed genome engineering. We present a modification to the TRACE approach which allows genetic manipulation to be mediated by homology directed repair (HDR) involving short (50 bp) regions of homology with only transient expression of Cas9 and sgRNA components. Furthermore, we demonstrate that increased efficiency can be achieved using a strain in which Cas9 is constitutively expressed by prior integration into the genome and the sgRNA is expressed transiently after electroporation of a PCR product. Finally, we describe use of a sequence that encodes codon-optimized mNeonGreen with a tandem purification tag for C-terminal tagging of genes. These tools enable short-homology genome engineering for the first time in *C. neoformans*.

## RESULTS

### Expression optimization of Cas9 for *C. neoformans*

We reasoned that improved Cas9 expression in *Cryptococcus neoformans* might allow for short homology (50 bp) mediated HDR. Codon optimization can increase gene expression, and introns are required for efficient expression in *C. neoformans* (Goebels *et al.* 2013). Accordingly, we engineered the *CAS9* coding sequence to use the most common codon for each amino acid (as determined by using all predicted open reading frames), and further introduced the single intron from *C. neoformans* gene *CNAG_05249* (Fig. 1A). We chose this intron because it is average in length for *C. neoformans* (Burke *et al.* 2018), and we further modified the sequence to have a consensus 5’ splice site (GTATGT) and 3’ splice site (CAG). This was placed into the codon optimized *CAS9* at an arbitrarily chosen position (after amino acid 348). The optimized *CAS9* ORF was placed under the control of the *TEF1* promoter and terminator; this promoter also contains an intron in its 5’ untranslated region. Lastly, we added a sequence encoding two SV40 nuclear localization signals to the C-terminus of the *CAS9* coding sequence. We refer to this gene as *C. neoformans*-optimized *CAS9* or *CnoCAS9*.

**Figure 1.**
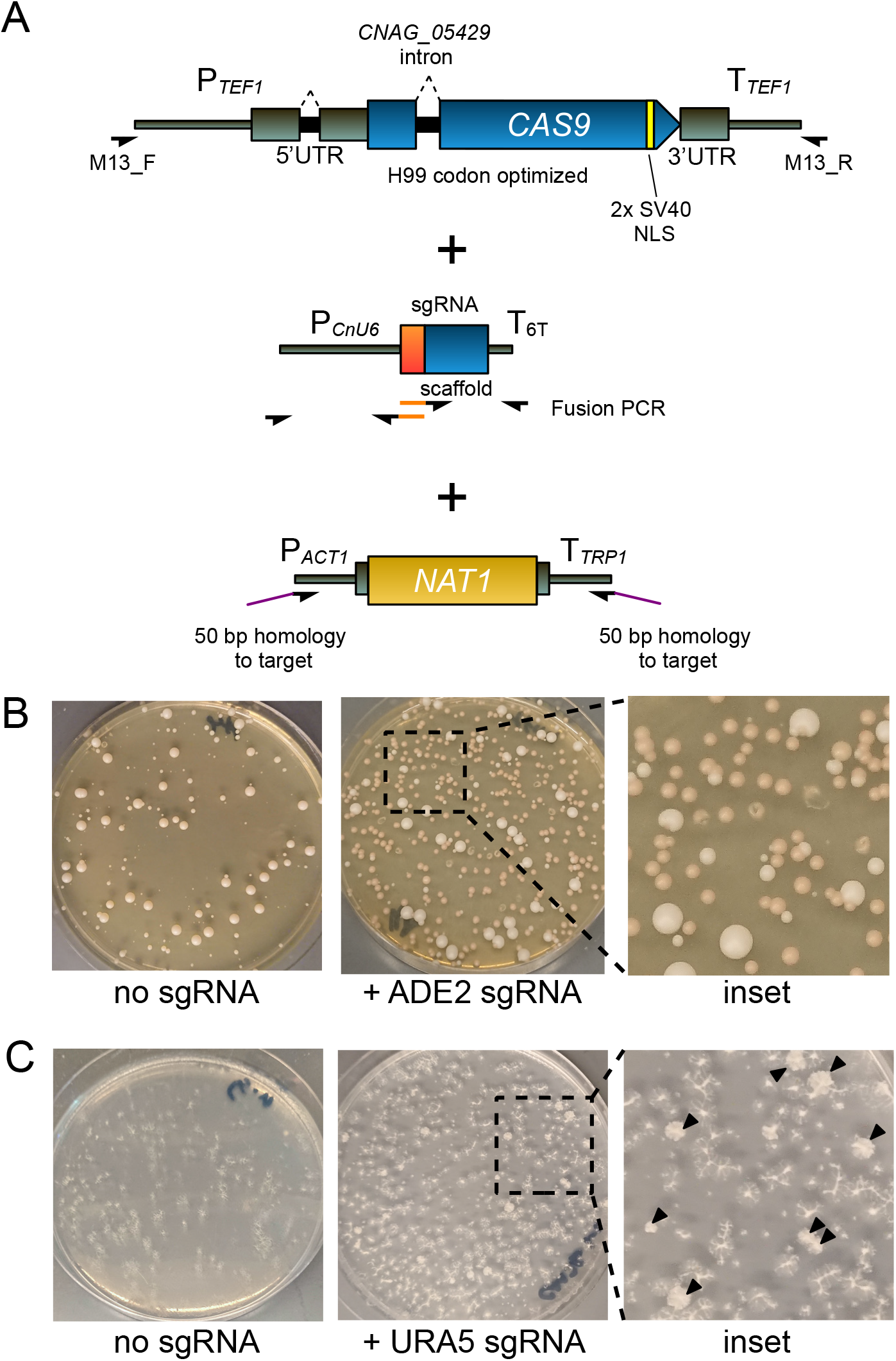
Optimized CRISPR enables short-homology editing in *C. neoformans*. (A) PCR scheme and schematic diagram for *CAS9* cassette with relevant optimizations, sgRNA cassette, and short homology deletion cassette. Orange segments denote 20 bp guide sequences introduced by fusion PCR. (B) Transformation plates for deletion of *ADE2* using transiently expressed optimized Cas9 and short homology deletion cassette. (C) Image of a *URA5* deletion transformation plate replica plated onto 5-FOA media. Black arrows indicate colonies growing on 5-FOA.

### *C. neoformans* optimized Cas9 enables short homology-driven homologous recombination

To test our strategy for short homology-based editing, we targeted the *ADE2* (CNAG_02294) locus, commonly used to assess the efficiency of gene editing techniques in fungi as the deletion strain accumulates a red pigment. We electroporated into KN99α linear DNAs corresponding to *CnoCAS9*, an *ADE2* sgRNA expressing cassette (*pU6-sgRNA*^*ADE2-1*^), and an *ade2*Δ::*NAT1* deletion cassette with 50 bp of homology on either end to the *ADE2* locus (amplified using ~70 bp oligonucleotides). No red colonies were detected when *pU6-sgRNA*^*ADE2-1*^ was omitted, whereas 69.4% of colonies from the electroporation that included *pU6-sgRNA*^*ADE2-1*^ showed red pigmentation (Table 1, Fig. 1B)

**Table 1.** Colony counts of Ade- and Ura- transformants

| Experiment: | Cas9 state | # aux - | # aux + | % aux - |
| --- | --- | --- | --- | --- |
| *ade2Δ* | transient | 174 | 76 | 69.6 |
|  |  | 158 | 69 | 69.6 |
|  |  | 129 | 58 | 69.0 |
| *ura5Δ* | transient | 11 | 131 | 7.7 |
|  |  | 11 | 101 | 9.8 |
|  |  | 24 | 108 | 18.2 |
| *ade2Δ* | integrated | 1730* | 4 | 99.8 |
|  |  | 1950* | 8 | 99.6 |
|  |  | 1520* | 7 | 99.5 |
| *ura5Δ* | integrated | 82 | 53 | 60.7 |
|  |  | 106 | 17 | 86.2 |
|  |  | 69 | 22 | 75.8 |

\*values extrapolated from colony counts of a 1:100 dilution of transformed cells

We next targeted *URA5* (CNAG_03196) gene using the same method. We transformed KN99α with linear DNAs corresponding to *CnoCAS9*, sgRNA, *ura5*Δ::*NAT1* deletion cassette (*pU6-sgRNA*^*URA5-1*^). Transformants were replica plated onto media containing 5-fluoroorotic acid (5-FOA) to screen for Ura-mutants (Boeke *et al.* 1984). Although the efficiency of deletion was lower than for *ADE2*, we still readily obtained the desired transformants, with 11.9% of colonies growing when replica plated on 5-FOA (Table 1, Fig. 1C).

Although these data demonstrate that our optimized Cas9 system can be used to disrupt genes of interest, it was unclear if the mutants arose by HDR through 50 bp homology. Previous attempts with short homology using TRACE only disrupted the target locus through insertion of the marker, presumably via nonhomologous end-joining (Fan and Lin 2018). We predicted three potential resolutions of a Cas9 induced double stranded break (DSB): recombination could occur at both ends to yield full deletion/replacement; non-homologous end joining (NHEJ) could occur at both ends resulting in an insertion mutant; or recombination could occur at one end, but NHEJ could occur at the other end, resulting in a partial deletion (Fig, 2A). To determine whether the Ura- and Ade-transformants were insertion or deletion mutants, we designed PCR primers to test whether the desired HDR event had occurred at each end. PCR products should only be amplified if insertion via NHEJ occurred (Figure 2A). We used this approach to assess transformants from three replicates each of transformations targeting *ADE2* and *URA5*: 16 red colonies from each *ade2*Δ transformation and all colonies from *ura5*Δ transformations that grew on 5-FOA plates. Half of the *ade2* mutants underwent HDR at either the 5’ or 3’ end of the *ade2* ORF, and 61% of the *ura5* mutants underwent HDR at either end(Table 2; representative colony PCR results are shown in Figure 2B). Among these transformants we identified multiple *ade2* and *ura5* mutants that produced no PCR products from either the 5’ or 3’ ends of the targeted ORF, consistent with a full ORF deletion (Fig, 2B). We selected eight such *ade2* mutants for further analysis by Sanger sequencing of the upstream and downstream regions of the putative *ade2*Δ*::NAT1* ORF (amplified by PCR using primers annealing 450 bp upstream or downstream of the recombinational junctions and within the *NAT1*. This sequencing showed that both junctions from all eight mutants were consistent with precise HDR (Fig. 2C). These data demonstrate that the *CnoCAS9* system allows for precise deletion using 50 bp homology arms.

**Table 2.**
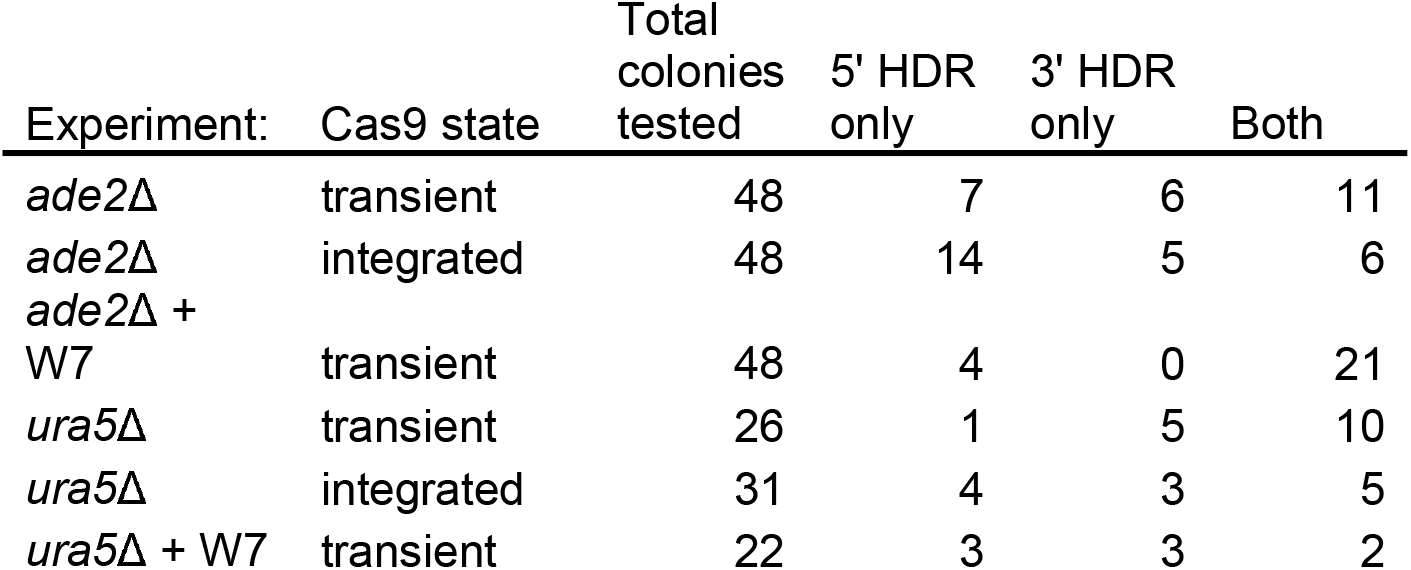
Colony PCR results by experiment

**Figure 2.**
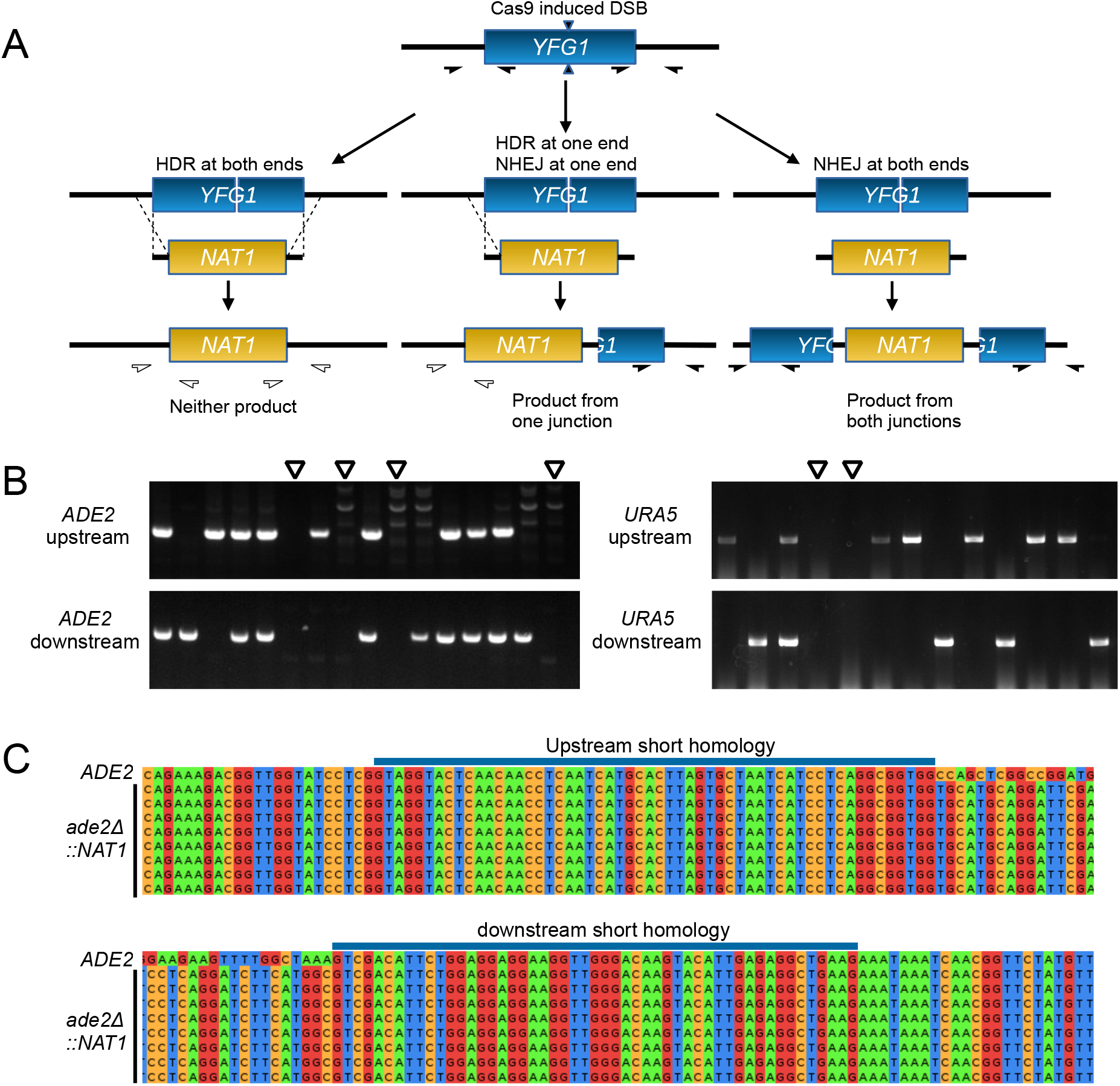
Genotyping of *ade2* and *ura5* transformants. (A) Possible outcomes of recombination outcomes and colony PCR genotyping scheme. Filled half-arrows indicate primer pairs that should amplify the target region. (B) Representative gels from colony PCR genotyping of transformants. Open arrowheads indicate candidates lacking both 5’ and 3’ junctions, consistent with the expected patterns for full *ade2*Δ and *ura5*Δ mutants. (C) Sanger sequencing of *ade2*Δ candidates from colony PCR genotyping.

Although 50 bp homology was sufficient to induce HDR, we frequently observed partial deletions with HDR at only one end. Arras and colleagues previously reported that supplementation of pre-culture media with W7 hydrochloride, an inhibitor of nonhomologous end-joining, increased the overall frequency of homologous recombination in the context of biolistic transformation (Arras and Fraser 2016). We reasoned that similar inhibition of NHEJ might result in increased HDR events. We tested this by adding 10 μg/ml W7 to both the preculture and outgrowth media for electroporation while targeting the *ADE2* locus for deletion as previously described. Colony PCR showed that while including W7 did not change overall transformation efficiency, it did increase the co-occurrence of HDR in the same transformant (Fig. S1). Without W7, 45% (11/24) of colonies that underwent HDR showed a full deletion, compared to 84% (21/25) with W7 supplementation (n=3, p = 0.015, Student’s t-test) (Table 2). We did not observe any change in HDR frequency for deletion of *URA5*. These data show that W7 supplementation may increase HDR, but that this effect maybe sgRNA- and/or locus-dependent.

### Stable integration of *CnoCAS9* increases transformation and homologous recombination efficiency

We observed low transformation efficiency when we attempted to generate a *URA5* deletion, despite testing multiple distinct sgRNA-encoding cassettes (data not shown). Fan and colleagues have reported that increasing the concentration of CAS9-encoding DNA or sgRNA-encoding construct improved both editing and transformation efficiency (Fan and Lin 2018) and Arras and colleagues previously found that their system required a genome-borne copy of the *CAS9* gene to function effectively (Arras *et al.* 2016). We reasoned that a genome-borne copy of *CAS9* might have greater expression than transiently expressed Cas9, and could thereby increase CRISPR-Cas9 editing efficiency. Additionally, this approach could reduce the number of separate DNA molecules that a given cell would need to take up during transformation, potentially increasing desired outcomes. Two loci have been reported to be ‘Safe Havens’ for integration of transgenes in *C. neoformans*, Safe Haven 1 (SH1) and Safe Haven 2 (SH2), where gene insertion does not impact fungal virulence (Arras *et al.* 2015; Upadhya *et al.* 2017). We inserted a *CnoCAS9* marked with *HYGB* at the *SH2* locus using TRACE without any flanking homology (See Methods) And confirmed integration by colony PCR of the *SH2* locus.

This resulting strain, CM2049 showed drastically increased editing efficiency (Figure 3). Transformation efficiency increased from approximately 300 colonies per ug DNA using transiently expressed Cas9 to more than 1×10^5^ colonies per ug DNA with CM2049. Furthermore, when transforming CM2049 to delete *ADE2*, nearly all transformant colonies showed red pigmentation (Fig. 3A). For *URA5* deletion in CM2049, 74.2% of transformants grew on media containing 5-FOA (Table 1). Colony PCR of Ade- and Ura- transformants showed that several transformants had undergone HDR at both ends (Fig. 3B). Across 3 replicates, 7.4% of all transformants from *URA5* deletion experiments underwent homologous recombination at both junctions to delete *URA5* using transiently-expressed *CnoCAS9*, compared to 30.8% using genome-borne Cas9.

**Figure 3.**
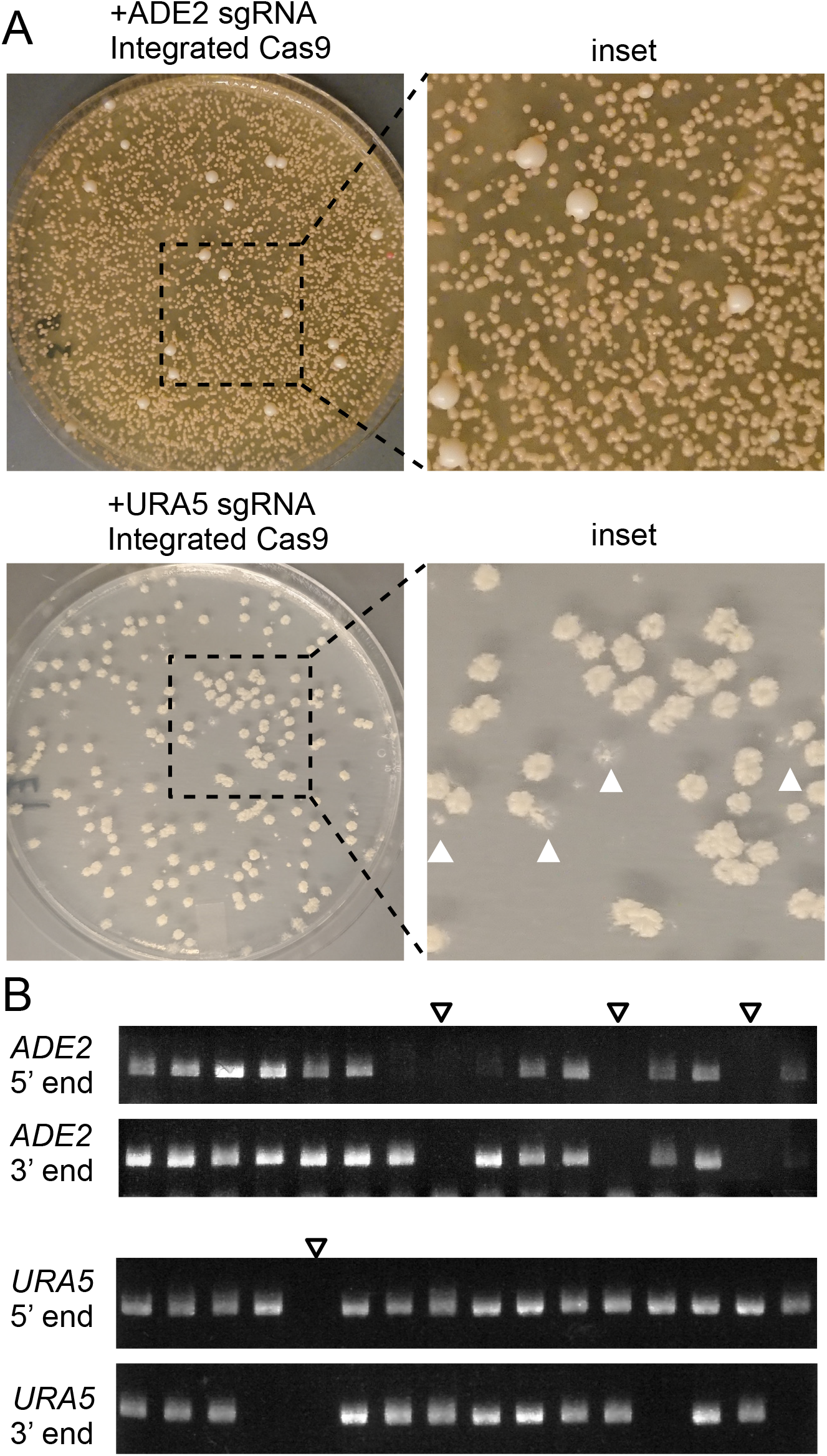
Deletion experiments using a genome-borne copy of optimized *CAS9*. (A) Transformation plates for deletion of *ADE2* or *URA5* in CM2049 using a short homology deletion cassette. The *URA5* plate shows transformants replica plated onto 5-FOA media. White arrowheads indicate examples of colonies that failed to grow on 5-FOA. (B) Representative gels from colony PCR genotyping of *ade2* and *ura5* transformants in CM2049. Empty arrowheads indicate candidates lacking both 5’ and 3’ junctions, consistent with the expected patterns for full *ade2*Δ and *ura5*Δ mutants.

### Integration of *CnoCAS* into the *C. neoformans* genome does not impact mammalian virulence

To test whether the strain harboring integrated *CnoCAS9* impacted infection and virulence, we inoculated groups of C57BL/6J mice intranasally (Cox *et al.* 2000) with the parental KN99α strain or the derivative expressing *CnoCAS9* (CM2049). Infection was allowed to proceed until mice reached their predefined experimental endpoint (see Methods). Survival curves of both groups of mice were similar (Fig. 4A) and there were no significant differences in lung or brain fungal burden at the time of sacrifice (Fig. 4B and C), indicating that integration of *CnoCAS9* did not influence fungal virulence or dissemination to the central nervous system (Fig. 4).

**Figure 4.**
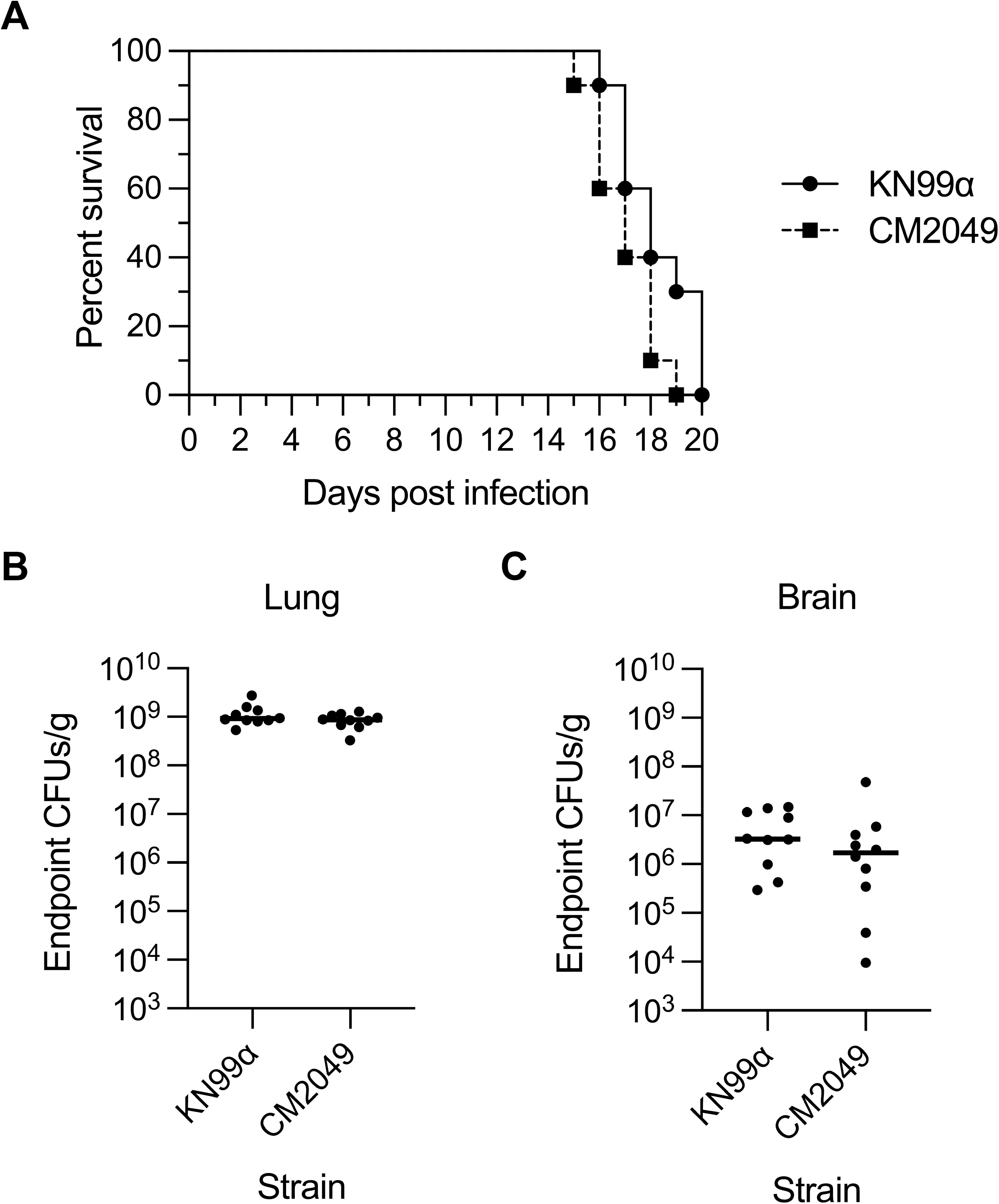
Evaluation of CM2049 infectivity in mice. Groups of 10 C57BL/6J mice were infected with 5 x 10^4^ cells of either CM2049 or its parent strain KN99α. (A) Survival curve. (B and C) CFUs per gram of organ tissue in the lungs (B) and brains (C) of mice at the time of sacrifice.

### Design and implementation of a localization and purification Tag

Using the codon optimization approach described above, we generated and tested optimized versions of several fluorescent proteins, including GFP, mCherry, and mNeonGreen. We found that a Cryptococcus-optimized mNeonGreen performed particularly well. We attached to this sequence a calmodulin-binding protein tag followed by 2X FLAG purification tag we have used in prior work (Dumesic *et al.* 2013; Dumesic *et al.* 2015a; Dumesic *et al.* 2015b; Burke *et al.* 2018; Catania *et al.* 2020; Summers *et al.* 2020) to produce a combined localization and purification (LAP) tag. We also appended a cleavage-polyadenylation/transcription termination sequence from the *CNAG_07988* gene and a *NAT1* selectable marker (Mcdade and Cox 2001). Amplification of this construct with 50 bp of targeting homology enabled us to design donor DNAs to C-terminally tag genes of interest. In each case, we also encoded an sgRNA targeting a region just downstream of the coding sequence. We transformed CM2049 with a donor construct produced by PCR, and a guide-expressing construct (Fig. 5). For one example, tagging the cryptococcal ortholog of the *S. cerevisiae* gene encoding the nucleoporin Nup107 (*CNAG_04149*), yielded a pattern consistent with the expected nuclear rim staining, and a protein product of the anticipated size of 132 kD (Fig. 5).

**Figure 5.**
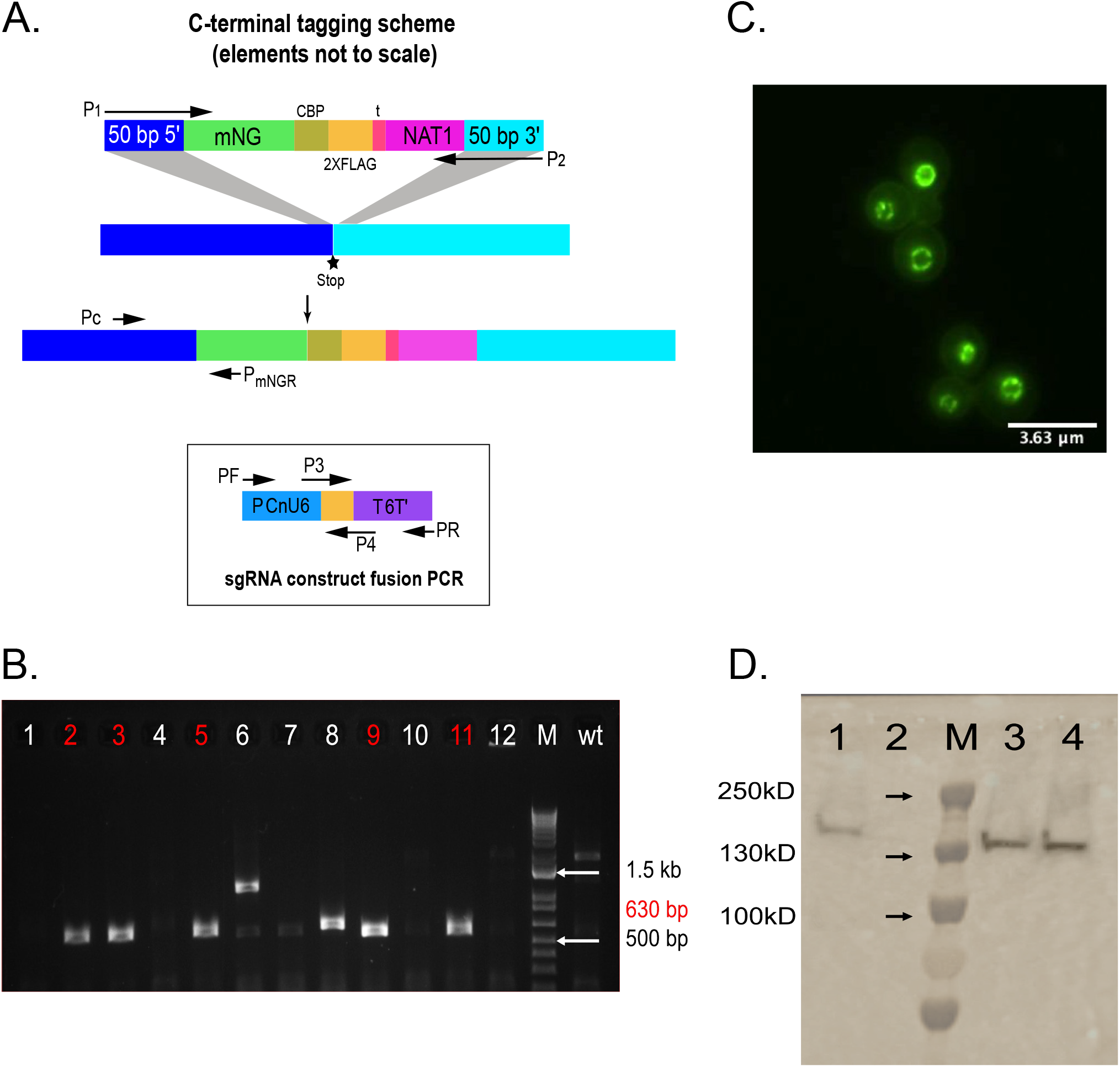
A localization and purification (LAP) tag for *C. neoformans*. (A) Scheme of the LAP-tagging strategy. (B) Colony PCR testing of 12 colonies and the parent strain to verify LAP tagging of *NUP107*. Positive colonies are highlighted in red. (C) Imaging of colony from panel B, using a 100X oil immersion objective and a FITC filter. (D) Immunoblot analysis. 1, positive control (*CCC1-CBP-2X-FLAG*, constructed by biolistic transformation); 2, parental strain; 3, *NUP107*-*mNeonGreen-CBP-2XFLAG* positive clone #1; 4, *NUP107-mNeonGreen-CBP-2XFLAG* positive clone #2.

To test our LAP tag in other cellular contexts, we tagged several additional genes selected based on information on their orthologs in *Saccharomyces cerevisiae* or studies of localization in *C. neoformans.* These genes encode the mitochondrial co-chaperone Mrj1 (Horianopoulos *et al.* 2020), the plasma membrane ATPase Pma1 (Farnoud *et al.* 2014), the zinc finger transcription factor Cqs2 (Tian *et al.* 2018; Summers *et al.* 2020) and an ortholog of the *S. cerevisiae* endoplasmic reticulum chaperone Erj5 (Carla Fama et al. 2007). All of these yielded the expected localization patterns (Fig. 6), suggesting that our tag will be a useful tool for studies of cryptococcal cell biology.

**Figure 6.**
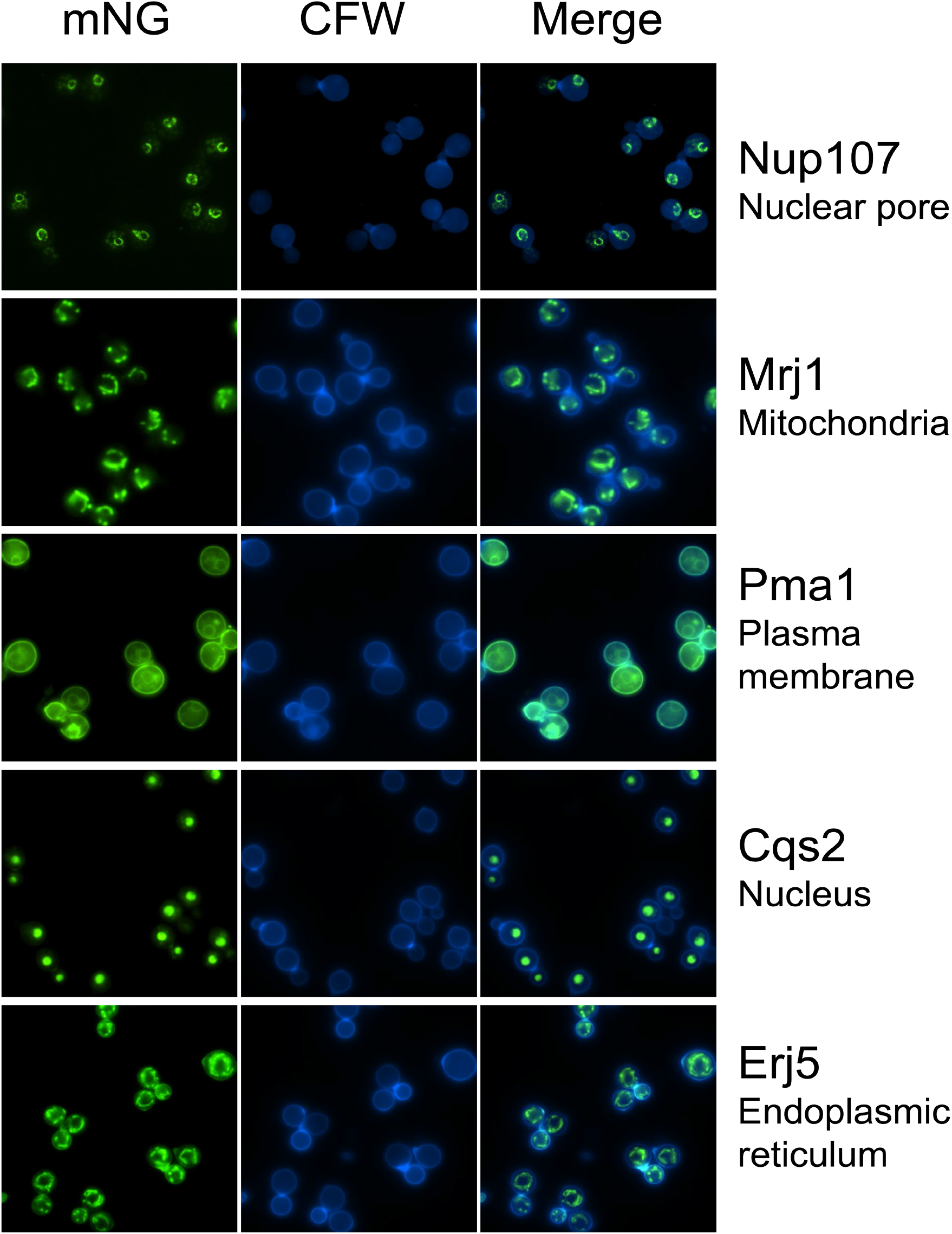
Proteins with distinct predicted localization patterns can be visualized with the *C. neoformans* LAP tag. Shown are representative images of strains expressing the indicated proteins tagged with mNeonGreen-CBP-2XFLAG; expected localization is included at the right. mNeonGreen fluorescence in the FITC channel is shown in the left column and calcofluor staining of cell wall chitin in the DAPI channel is shown in the center column.

### Testing of portability and reproducibility

The tools described were developed in the Madhani laboratory. We sought to verify that our methods could be deployed in a different laboratory setting. We provided protocols and the *CnoCAS9* expressing strain *CM2049* to the Doering group, who successfully deleted *CNAG_01050* and tagged *NUP107* using these methods (Fig. S2). These data suggest that the tools described here are robust.

## DISCUSSION

Short homology-driven genome modification, first developed in *S. cerevisiae* (Baudin et al. 1993), enables rapid strain engineering, and was pivotal to the construction of whole-genome knockout and tag collections in *S. cerevisiae* (Giaever *et al.* 2002; Ghaemmaghami *et al.* 2003; Huh *et al.* 2003; Giaever *et al.* 2004). Cloning or fusion PCR to join long homology arms to a marker or construct of interest can be difficult and require extensive troubleshooting. Furthermore, a custom construct must be cloned for every desired manipulation, frustrating efforts to study multiple genes in parallel. Although CRISPR-Cas9 methods have been developed for use in *C. neoformans*, these methods have not addressed this major bottleneck.

In this work, we report a series of modifications to *CAS9* to optimize Cas9 expression in *C. neoformans*. We show that the optimized *CAS9* gene successfully allows use of short homology for genetic manipulation, results that have been replicated by a second group. Our optimized version represents an improvement over current *CAS9* genes used in *Cryptococcus* research and, consistent with previous observations by Arras and colleagues for the human-optimized Cas9 (Arras *et al.* 2016), integration of this protein does not alter fungal virulence. Although our construct was optimized specifically for *C. neoformans*, we expect that it may be of broader utility, given the previously observed activity of *Caenorhabditis elegans* codon optimized Cas9 activity in other *Cryptococcus* species (Fan and Lin 2018).

Another approach to improving short homology-based editing in species that display high levels of NHEJ is the incorporation of single stranded DNA strategies. Single stranded DNA oligonucleotides (ssODN) were first shown to increase editing efficiency when using zinc-finger nucleases in human cell lines (Chen *et al.* 2011). With long or short ssODNs, editing in induced pluripotent stem cells, rats, and zebrafish has been achieved using 30, 60, and 40 bp homology arms respectively (Bai *et al.* 2015; Yoshimi *et al.* 2016; Okamoto *et al.* 2019). An efficient 100 bp homology arm knock-in strategy for *Drosophila* has been described that generated long ssDNA cassettes by enzymatic digestion of a PCR product with one 5’ phosphorylated strand (Kanca *et al.* 2019). Likewise, a 120 bp single stranded homology arms on a hybrid donor can produce efficient editing in *C. elegans* (Dokshin *et al.* 2018). Although these approaches were not required for successful short homology editing in *C. neoformans*, they may allow further increases in efficiency. Initial efforts to transform KN99α with ssDNA by electroporation were not successful (data not shown), but further optimizations may open this promising avenue.

## METHODS

### Strains and Media

All strains were maintained in 20% glycerol stocks stored at −70°C. *C. neoformans* and *Saccharomyces cerevisiae* strains were recovered from frozen stocks on YPD media (2% Bacto Peptone, 2% dextrose, 1% yeast extract) with agar for 2-3 days at 30°C or directly in liquid YPD media overnight at 30°C in a roller drum incubator at 60 RPM. Precultures for yeast transformation and *C. neoformans* electroporation, as well as outgrowths to allow for recovery following electroporation were in liquid YPD media. Following electroporation, transformants were selected on YPD plates supplemented with either 125 μg/ml nourseothricin or 300 μg/ml hygromycin as appropriate. Screening for Ura- transformants was performed on solid YNB media (0.67% Difco yeast nitrogen base without amino acids, 2% dextrose, 2% agar for solid media) supplemented with 1 mg/ml 5-fluoroorotic acid (5-FOA) and necessary amino acids. For plasmid cloning experiments using gap repair in *S. cerevisiae*, transformants were selected on YNB media lacking uracil.

A list of strains, primers, and plasmids used in this study is provided in Supplementary Table 1.

### Plasmid Construction

Cas9 was optimized based on codon frequency in *C. neoformans* var. *grubii* strain H99. Specifically, we generated a codon table for each amino acid using all annotated H99 coding sequences. We identified the most common codon for each amino acid. Additionally, we designed the codon optimized Cas9 gene to contain an intron from CNAG_05429. This intron was modified to have consensus 5’ and 3’ splice sites. This entire *C. neoformans*-optimized (Cno) Cas9 construct was ordered from GenScript (Piscataway, NJ) through their gene synthesis service and was received as an insert in pUC57 flanked by XbaI and BamHI restriction sites. The pUC57-Cas9 plasmid was digested with XbaI and BamHI to liberate the *CnoCAS9*. The *TEF1* promoter and terminator were amplified by PCR from H99 genomic DNA using primers P01 and P02 for the promoter, and P03 and P04 for the terminator, containing homology to the pRS316 multicloning site and *CnoCAS9*. The *TEF1* promoter, optimized Cas9 construct, and *TEF1* terminator were cloned into pRS316 digested with XbaI and HindIII by gap repair in *S. cerevisiae* strain BY4741. *S. cerevisiae* transformations were performed by the lithium acetate procedure as described elsewhere (Gietz and Schiestl 2007). pRS316 DNA was recovered from resulting transformants using a Zymoprep Yeast Plasmid Miniprep Kit (Zymo Research, Irvine, CA) and electroporated into *Escherichia coli* strain DH5α to isolate single plasmid copies. Plasmid was recovered from candidate *E. coli* using a NucleoSpin Plasmid, Mini kit (Macherey-Nagel, Düren, Germany) and verified by Sanger sequencing (Genewiz, South Plainfield, NJ) using custom oligos P05-P12. One resulting correct clone was designated pBHM2403.

The *HYGB* marker was inserted upstream of *CAS9* in pBHM2403 to yield pBHM2408. Briefly, *HYGB* was amplified from pBHM2402 (S. Catania, unpublished data) using primers P13 and P14 containing homology to the NotI and SacII sites of pRS316/pBHM2403. The *HYGB* marker on pBHM2402 contains the same *ACT1* and *TRP1* promoter and terminator sequences as the *NAT1* marker. This PCR product was cloned into pBHM2403digested with NotI and SacII by Gibson assembly using the NEB Hifi DNA Assembly Kit (New England Biolabs, Ipswich, MA). To generate a sgRNA expression cassette, the *Cryptococcus* native U6 promoter was amplified from JEC21 genomic DNA (Wang *et al.* 2016) using P15 and P16, containing homology to the sgRNA scaffold. The sgRNA scaffold followed by 6T terminator were amplified from pSDMA66 (Arras *et al.* 2016) using P17, containing homology to the U6 promoter, and P18. These PCR products were cloned by Gibson assembly into pRS316 digested with BamHI and HindIII. Candidate colonies were Sanger sequenced using universal primers M13_F and M13_R and an identified correct clone was designated pBHM2329.

The tagging construct template plasmid was generated by homologous recombination in yeast strain BY4741. Codon-optimized mNeonGreen, CBP-2X FLAG-UTR and *NAT1*/*HYGB* segments were amplified from independent plasmids using primers P19 to P24 and transformed alongside pRS316 vector digested with NotI and ClaI. Yeast transformation was done using lithium acetate protocol followed by extraction of pRS316 plasmid from yeast cells as described (Hoffman and Winston 1987). Plasmid was isolated by transformation of DH5α by electroporation. Individual colonies were used for isolating plasmid and clones were verified by enzymatic digestion and gel electrophoresis followed by Sanger sequencing. Correct plasmids were designated as pBHM2404(*NAT1*) and pBHM2406(*HYGB*).

### Electroporation

Electroporations were performed as described by Lin and colleagues with the following modifications (Fan and Lin 2018). Briefly, cultures were inoculated from overnight cultures into 100 ml YPD at an OD_600_ of 0.2 and grown for 4-5 hours until an OD_600_ of 0.65-0.8 was reached. Cells were collected by centrifugation and washed twice in ice cold water, before being resuspended in electroporation buffer (10 mM Tris-HCl pH 7.5, 1 mM MgCl_2_, 270 mM sucrose) (Fan and Lin 2018) and incubated at 4°C with 1 mM DTT for one hour. Cells were collected and resuspended in 250 μl of fresh ice-cold electroporation buffer for an approximate resulting cell-buffer slurry volume of 450-500 μl. 50 μl of cells were mixed with PCR products and transferred to a precooled 2 mm gap electroporation cuvette (Bio-Rad Laboratories, Hercules, CA). A BTX Gemini X2 instrument was used for electroporation with the following settings: 500 V, 400 Ω, 250 μF. Cells were resuspended following electroporation in 1 ml YPAD and incubated at 30°C for 1 hour before plating on selective media.

### Primers, PCR products for deletion experiments

Primers were ordered from IDT using their 25 nmole DNA Oligo or 100 nmole DNA Oligo service as appropriate. Deletion cassettes for *URA5* and *ADE2* were amplified from pCH233 using primer pairs P25+P26, and P27+P28, respectively. Primer pairs contained approximately 50 bp of homology to the respective gene of interest. PCR was performed with ExTaq (Takara Bio Inc., Kusatsu, Shiga, Japan) per manufacturer’s instructions, supplemented with 2% DMSO. For transient Cas9 experiments, the *CAS9* cassette was amplified from pBHM2403 using ExTaq per manufacturer’s instructions and primers M13_F and M13_R.

Cassettes expressing sgRNAs were amplified in two PCR steps. Briefly, a set of primer pairs containing the 20 bp target sequence was designed to amplify the U6 promoter and sgRNA scaffold with 6T terminator, respectively, from pBHM2329. U6 and scaffold products were then mixed in equal volumes and used as the template for fusion PCR using a cocktail of Pfu and Taq polymerases as previously described (Chun and Madhani 2010). For example, to produce the *ADE2-1* sgRNA, the U6 promoter was amplified from pBHM2329 using M13_F andP30. The sgRNA scaffold and 6T were then amplified from pBHM2329 using P29 and M13_R. The two products were then joined by fusion PCR using primers P33 and P34. The URA5 sgRNA was produced similarly using primers P31 and P32.

The sizes of PCR products were verified by gel electrophoresis. All PCR products were purified using a Macherey-Nagel Gel and PCR clean-up kit following their PCR cleanup procedure and eluted in a small quantity of sterile ddH2O. Each PCR aliquot of 100 μL was cleaned in an individual column and eluted in 15-20 μL sterile ddH2O to ensure high DNA concentration.

### Primers, PCR products for tagging experiments

Plasmid pBHM2404 (*NAT1*) was used as template for donor PCRs using primers with ~50 bp homology to the genome sequence (primers P51 through P60). Each PCR product was verified by gel electrophoresis and purified using Macherey-Nagel Gel and PCR clean-up kit as described above. PCR products specifying sgRNA cassettes targeting downstream regions of each gene were also produced as described above, using primers P61 through P70.

### Strain Construction

To delete *URA5* or *ADE2*, KN99α (CM026) was electroporated with 700ng of *URA5* or *ADE2* sgRNA, 1 μg of *CAS9* cassette, and 2 μg of the *URA5* or *ADE2* deletion cassette, as appropriate. CM2049 was constructed by electroporating KN99α (CM026) with 700ng of *SH2* sgRNA and 2 μg *HYGB-CAS9* amplified from pBHM2408 using primers M13_F and M13_R. *HYGB-CAS9* was amplified using Q5 High-Fidelity DNA Polymerase (New England Biolabs) per manufacturer’s protocols. SH2 sgRNA was produced as described earlier using primers P47 and P48. Insertion of *HYGB-CAS9* at the SH2 locus was verified by colony PCR using primers P49 and P50.

To tag the C termini of genes of interest, CM2049 was electroporated with 700ng of CNAG specific sgRNA and 2μg of mNeonGreen-CBP-2xFLAG donor PCR product with CNAG specific 50 bp homology arms.Transformants were selected on YPAD + NAT plates.

### Colony PCR

A detailed stepwise protocol with images for Colony PCR is available in Supplementary File S1. Briefly, cells were patched from a transformant colony onto fresh selective media. Next, a small (1-2 ul) quantity of cells was smeared evenly against the bottom of a PCR tube using a wooden toothpick. The cells were microwaved for 2 minutes, and PCR master mix was immediately added to the lysed cells. Colony PCR bands were assessed by gel electrophoresis in a 1% agarose gel.

Primers P39 + P41, P40 + P42 were used for PCR genotyping of the *ade2* locus. Primers P44 + P45, P43 + P46 were used for PCR genotyping of the *ura5* locus. Primers P71 + P72 through P76 were used for genotyping of their respective CNAG C-terminal tag candidates.

### Genomic DNA purification and Sanger sequencing

Genomic DNA was purified by CTAB extraction. Briefly, cells from a 5 ml overnight culture were collected by centrifugation. Cells were washed once in sterile distilled water, following which the supernatant was entirely discarded and the pellet was frozen in liquid nitrogen or at −80°C. 5 ml CTAB buffer (100mM Tris pH 7.4, 0.7M NaCl, 10mM EDTA pH 8.0, 1% CTAB (Sigma-Aldrich, St. Louis, MO)) with 1% β-mercaptoethanol was prewarmed to 65°C and then added to the frozen pellet. The pellet was then incubated for 2 hours at 65°C with periodic agitation to lyse the cells. DNA was then isolated by one or two rounds of chloroform extraction as necessary followed by isopropanol precipitation. The pellet following precipitation was resuspended in 400 ul TE buffer and treated with 1 ul of 10mg/mL RNAse A with incubation at 37°C for 1 hour. RNAse treated samples were then treated with 5 ul 20 mg/mL Proteinase K with incubation at 55°C for 1 hour. DNA was then isolated again by phenol:chloroform extraction followed by ethanol precipitation. Pellets were resuspended in 100 ul distilled water.

Upstream and downstream junctions for the *ADE2* ORF were amplified using primers P35 and P37, P36 and P38 respectively. PCR products were sent to Genewiz for Sanger sequencing using primers P41 or P42.

### Murine infection model

All work was performed under an approved institutional animal protocol. *C. neoformans* strains KN99α (CM026) and CM2049 were grown overnight in YPAD medium, washed twice in sterile saline, counted by hemocytometer, and resuspended to a concentration of 1 x 10^6^ cells/ml in sterile saline. Groups of 6-12-week-old C57BL/6J mice were anesthetized by intraperitoneal injection of 75 mg/kg ketamine and 0.5 mg/kg dexmedetomidine. Mice were suspended by their front incisors from a silk thread, and 50 μl of yeast suspension (5 x 10^4^ total yeast) were slowly pipetted into the nares. Mice were kept suspended for 10 minutes post inoculation, after which they were lowered, and anesthesia was reversed by intraperitoneal injection of 1 mg/kg atipamezole. Mice were monitored daily and, upon loss of 20% body weight relative to pre-infection or other signs of severe disease were euthanized by CO_2_ inhalation followed by cervical dislocation. Lung and brain tissue were harvested, resuspended to 5 ml final volume in sterile PBS, homogenized, and plated onto YPAD agar to determine CFUs.

### Fluorescence Microscopy

#### Sample preparation

5ml YNB media supplemented with 2% glucose was inoculated with each tagged strain followed by incubation at 30°C.Log stage cultures were used for imaging. 200 μl of culture suspension was centrifuged at maximum speed and the pellet was washed once with 200 μl PBS and pelleted again at maximum speed. The cell pellet was then resuspended in PBS with 100 μg/ml calcofluor white and incubated for 10-15 min in the dark at room temperature. Cells were pelleted again and washed once with PBS and once with sterile water. Finally, the cell pellet was resuspended in 50 μl sterile water and 10 μl of the cell suspension was spotted on glass slide and covered with coverslip for imaging.

#### Microscopy and image processing

Images were collected using Nikon ECLIPSE Ti2 microscope using an 100X oil immersion objective. Images were processed for background subtraction and color correction using NIS elements and ImageJ (Schneider *et al.* 2012).

### Protein extraction and immunoblotting

Protein extraction was done using previously described protocols (Catania *et al.* 2020). In brief, two milliliters of culture at OD_600_ = 1 was collected by centrifugation, and the cell pellet was frozen in liquid nitrogen. The cell pellet was then resuspended in 200 μl of 10% TCA and incubated on ice for 10 min. Cells were collected by centrifugation and washed once with 100% acetone and air-dried for 10 min. The pellet was then resuspended in 80 μl 1M Tris (pH 8.0) and 200 μl 2x Lysis buffer (Genescript LDS sample buffer 4X) followed by addition of glass beads. Sample was then boiled for 5 min at 100°C followed by bead-beating twice for 90 s. The cell lysate was centrifuged to remove residual cells. 25 μl of sample was subjected to SDS-PAGE (Genescript SurePAGE™, Bis-Tris gel) and transferred to a nitrocellulose membrane. Blotting was performed using anti-FLAG M2 antibody (1:3000, cat# F3165, Sigma) in 5% milk in TBS-T (10 mM Tris-HCl pH 7.6, 150 mM NaCl, 0.1% Tween 20) at 4°C overnight followed by three washes of 10 min each in TBS-T. The membrane was then incubated with anti-rabbit antibody conjugated to HRP (Bio-Rad) (1:8000 in 5% milk+TBS-T) for 60 min at room temperature followed by three washes,10 min each in TBS-T. The membrane was then incubated in SuperSignal West Pico reagent (Thermo Scientific) for 5 min. and visualized using Azure c400 instrument (Azure Biosystems).

## Data Availability

Plasmids pBHM2403, pBHM2404, pBHM2406, and pBHM2329 are available through Addgene.

## ACKNOWLEDGEMENTS

This work was supported by National Institutes of Health grants T32HL007185 (M.J.B.), F32AI152270 (M.J.B.), F31AI150194 (L.C.L), T32GM007067 (E.A.G.), R01AI087794 and R01AI000272 (H.D.M.)

---

**Figure S1.**
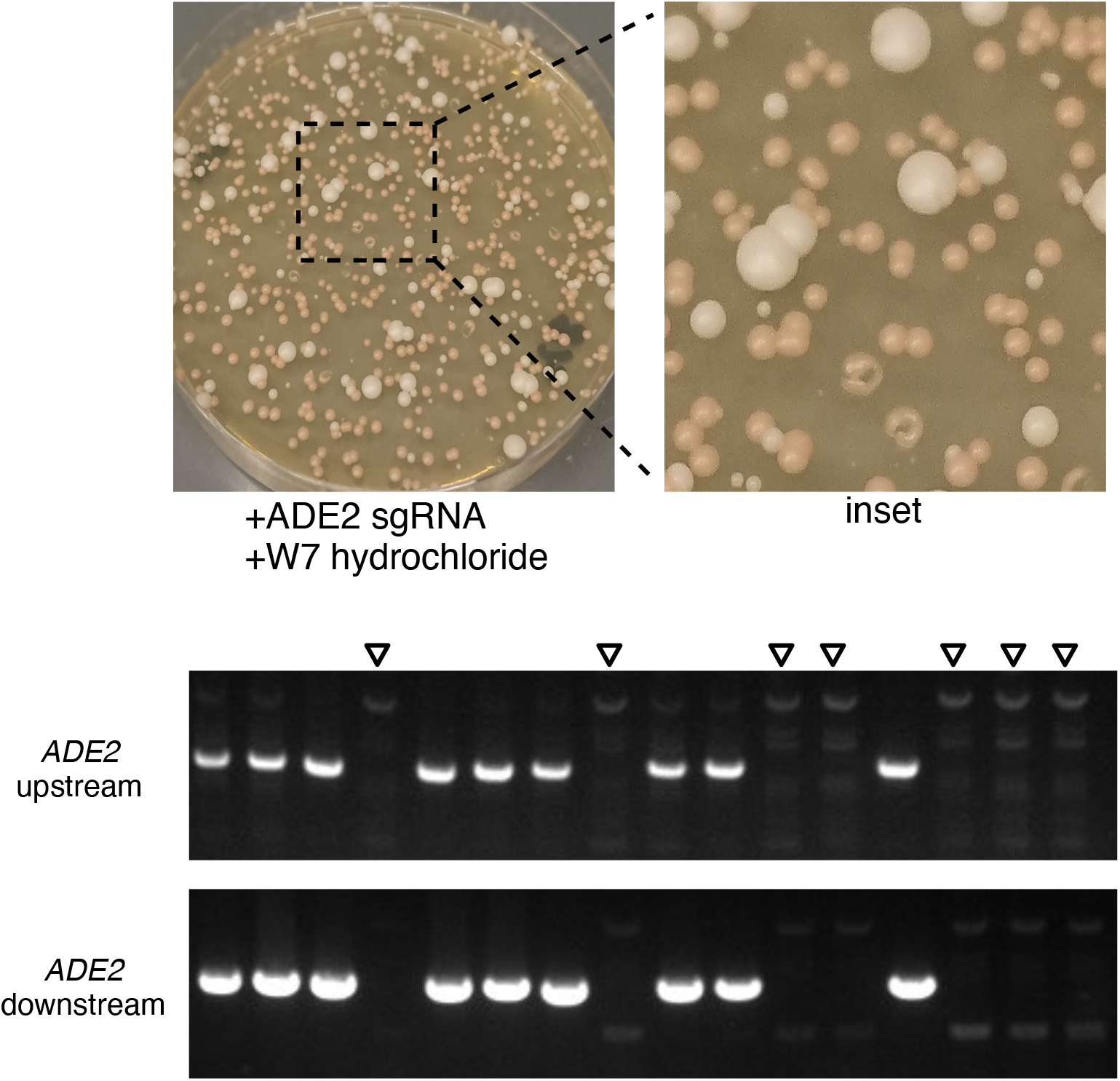
Use of W7-hydrochloride in deletion of *ADE2*. Transformation plate showing pink colonies consistent with ade2 disruption and colony PCR genotyping gel for 16 colonies are shown. Empty arrows indicate candidates lacking 5’ and 3’ junctions, consistent with expected patterns from full *ade2*Δ mutants.

**Figure S2.**
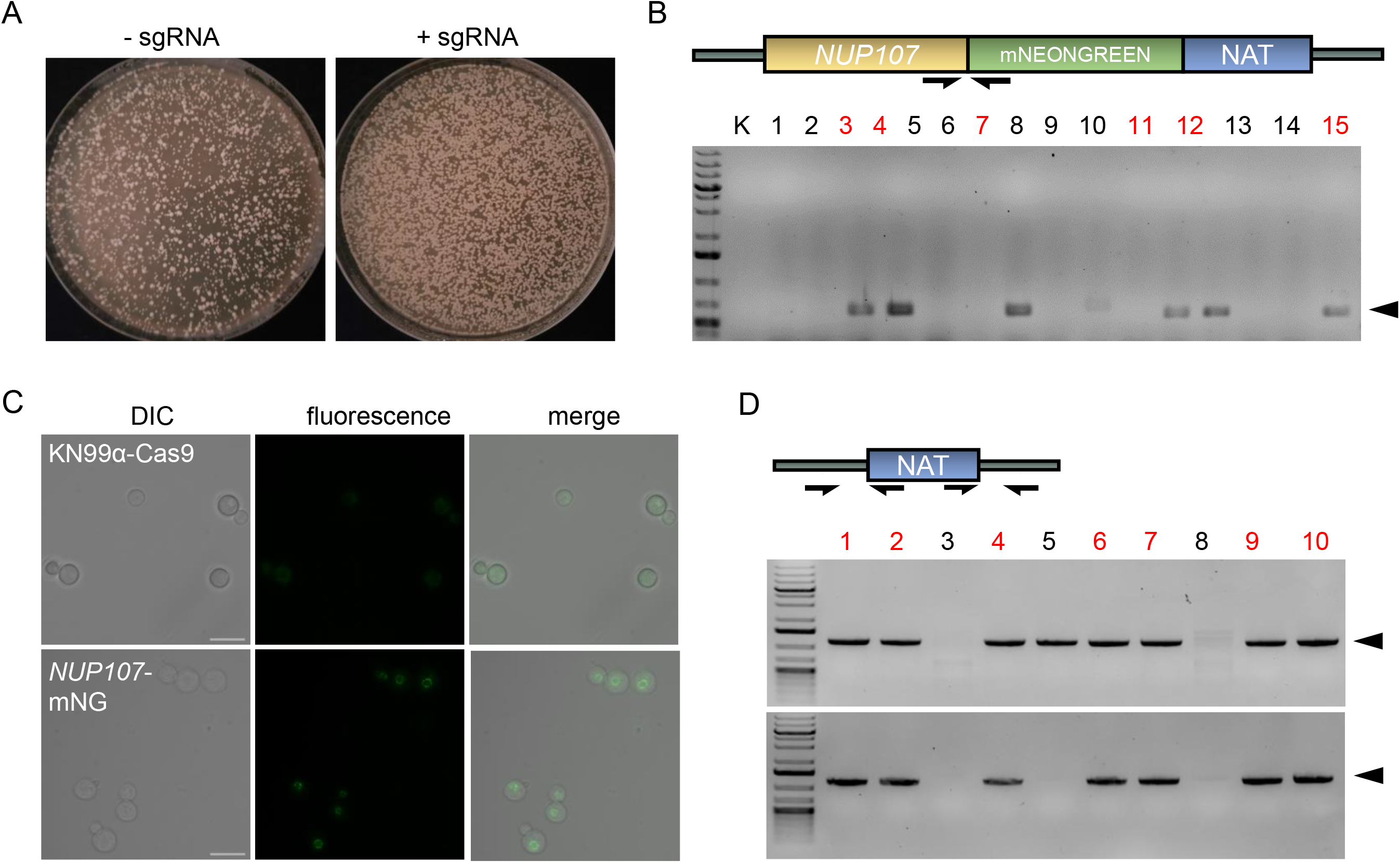
Successful application of optimized CRISPR methods by an independent group. (A-C) Use of the reported methods for modification of *NUP107* to encode a C-terminal mNeonGreen-CBP-2xFLAG tag. (A) Colony growth on nourseothricin-containing medium after transformation in the absence (left) or presence (right) of sgRNA. (B) Genotyping strategy and colony PCR assessment of the parent strain and 15 randomly selected transformants. Arrowhead, expected product size (630 bp); red text, verified transformants. (C) Transformant 12 imaged by DIC and fluorescence microscopy (using 100X oil objective and EGFP filter). Scale bar, 10 μm. (D) Use of the reported methods to delete CNAG_01050. Shown are the genotyping strategy and colony PCR assessment of 10 randomly selected transformants. Upper gel, 5’ primer pair (arrowhead marks the expected product size of 1167 bp); lower gel, 3’ primer pair (arrowhead marks the expected product size of 1123 bp); red text, verified transformants. The outer primer in each pair is beyond the region of homology used to mediate recombination.

