## Supplementary File S1 for "Short homology-directed repair using optimized Cas9 in the pathogen *Cryptococcus neoformans* enables rapid gene deletion and tagging"

*C. neoformans* Colony PCR Protocol Apr. 2021

Day 1

1. Patch colonies from transformation plate or other source onto fresh YPD plates with appropriate selection.

Day 2

2. Prepare colony PCR master mix as follows, mix by pipetting, and place on ice:

| | |
|---|---|
| 10x Pfu Buffer | 2.5 ul |
| 2.5 mM dNTPs | 2.5 ul |
| 100% DMSO | 1.25 ul |
| Primer-F, 10uM | 1.0 ul |
| Primer-R, 10uM | 1.0 ul |
| 1x Pfu enzyme | 0.5 ul |
| 5x Taq enzyme | 0.5 ul |
| dd $H_2O$ | 16.5 ul |

3. Using a sterile wooden toothpick, pick a small quantity (1-2 ul) of cells from a fresh patch.

   *Note: patches must be less than 48 hours old. Preferably <24 hours.*

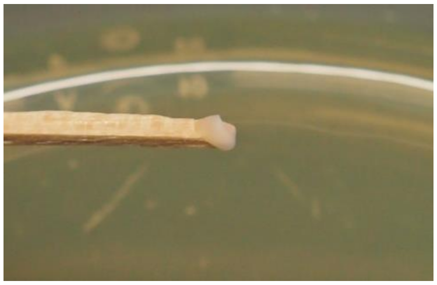

4. Smear cells against bottom of PCR tube in a thin even layer.

   *Note: Smear by pressing the shaft of the toothpick near the tip against the tube wall.*

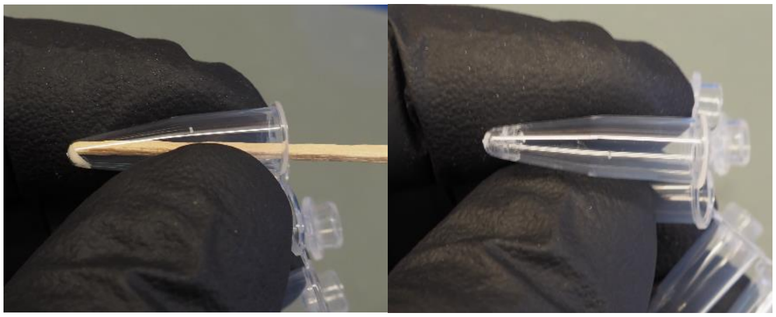

5. Repeat for all patches. Close lid of PCR tube and place in empty P10 tip rack.

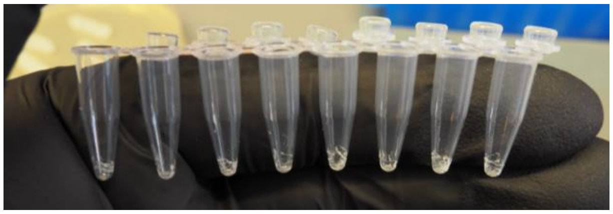

6. Lyse cells in microwave for 2 minutes.

   *Note: Microwave used in the lab is a 1200W Panasonic countertop microwave. Cells may appear slightly translucent after microwave lysis.*

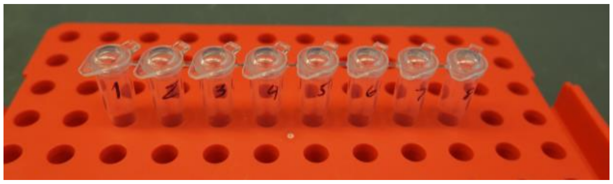

7. Pipette 25 ul of PCR master mix into each PCR tube. Keep PCR tubes on ice.

8. Vortex briefly to mix.

9. Run reaction in thermocycler (35 cycles, 95°C melting, 50°C annealing, and 72°C extension) as appropriate.

Additional Notes:

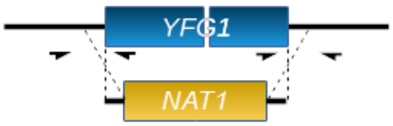


PCR primers should be designed to detect both the upstream and downstream recombinational junctions (e.g. [upstream forward, internal reverse], [internal forward, downstream reverse]).

Primers should lie outside of any homology regions. The length of the PCR product should be ideally less than 500 bp, and preferentially greater than 200 bp.

Colony PCR has also worked using 96 well PCR plates (Bio-Rad, Cat# HSP9601)

Materials:

10x Pfu buffer: 200 mMTris–HCl (pH 8.8), 20 mMMgSO4, 100 mMKCl, 100 mM(NH4)2SO4,1% (v/v) Triton X-100, 0.1% (w/v) BSA

2.5 mM dNTPs

100% DMSO

Colony PCR primers, 10 uM

Pfu, Taq polymerase

Sterile wooden toothpicks

PCR tube 8-strip (Greiner Bio-One Cat# 673283)

Thermocycler

1200W Microwave
